# The orbitofrontal cortex constructs allocentric schemas by integrating dynamic mobile agents with static environmental anchors

**DOI:** 10.64898/2026.03.01.708836

**Authors:** Ziyan Zhu, Bo Zhang, Xinyi Zhang, Yuji Naya

**Affiliations:** School of Psychological and Cognitive Sciences, Peking University, Beijing, China; Division of Humanities and Social Sciences, California Institute of Technology, Pasadena, United States; IDG/McGovern Institute for Brain Research at Peking University, Beijing, China; Beijing Key Laboratory of Behavior and Mental Health, Peking University, Beijing, China; Department of Psychiatry, Yale School of Medicine, New Haven, United States

**Keywords:** spatial schema, spatial layout, allocentric, egocentric, orbitofrontal cortex, lateral superior occipital cortex, hippocampus

## Abstract

Scene construction involves the automatic synthesis of fragmented visual inputs into coherent mental models. While previous work has focused on static landscapes, real-world scenes require integrating dynamic agents with stable environmental anchors. We investigated the neural hierarchy underlying this transformation using an automatic space-encoding task in a 3D virtual-reality environment and fMRI representational similarity analysis. Results revealed functional segregation across visual cortex: the superior lateral occipital cortex encoded the first-person perspective of mobile character layouts, whereas the lingual gyrus and occipital fusiform cortex represented environments specified by visible landmarks. The orbitofrontal cortex integrated these streams into an allocentric spatial schema, capturing relational geometry beyond the current visual field. Following encoding, these representations were expressed in the anterior hippocampus for self-localization. Together, the findings demonstrate an automatic hierarchical transformation in which the OFC serves as a central scaffold for constructing allocentric spatial frameworks underlying stable scene representation.

## 1. Introduction

Human vision provides only a narrow, fragmented window into the world, yet we perceive our surroundings as a seamless and stable whole^[1, 2]^. This experience relies on scene construction—the automatic process of synthesizing immediate visual inputs into a coherent mental representation ^[3, 4]^. Central to this process is the transformation of first-person perspective snapshots into allocentric spatial representations that define the global environment ^[5]^. While traditional research has focused on the perception of static landscapes ^[6, 7]^, real-world scenes are often populated by mobile objects, such as other people ^[4]^. However, it remains poorly understood how the dynamic spatial relationships among mobile objects are integrated with the environment specified by immobile landmarks to construct a coherent global scene.

Before a global scene can be constructed, the brain must first decompose the visible environment into its constituent spatial elements ^[8–12]^. This decomposition occurs within specialized visual areas that represent the world in egocentric, or viewpoint-dependent, coordinates. One essential stream involves representing the precise arrangement of dynamic agents relative to the observer ^[4, 13]^. Simultaneously, the brain must identify the immobile features that define the current environment, including both visible and non-visible landmarks ^[8, 10, 11, 14]^. By segregating mobile and immobile elements into distinct first-person representations, the visual system provides the raw material necessary to map immediate perceptions onto a coherent spatial environment that extends beyond the current visual field ^[15–17]^.

The integration of decomposed first-person streams into a unified allocentric spatial schema represents a higher-order stage of scene construction. Unlike immediate visual "snapshots" ^[18, 19]^, a schema is an abstract framework that captures the invariant relational geometry between mobile agents and the static environment ^[20]^. The orbitofrontal cortex (OFC) has emerged as a critical hub for this synthesis, encoding "latent" mental models that enable the brain to maintain a stable representation of the entire scene, even when key components move or are no longer directly visible ^[21, 22]^. This prefrontal schema provides the structural scaffolding that the hippocampus eventually utilizes to anchor the self within the global environment ^[4, 23]^.

A central question is whether this hierarchical process—from the decomposition of mobile and immobile elements to the construction of an allocentric schema—operates automatically during incidental perception. We developed the Automatic Space-Encoding (ASE) task to test whether the brain forms these high-dimensional models even when explicit spatial memorization is not required. Using Representational Similarity Analysis (RSA), we tracked how the first-person spatial layouts of human characters and landmarks are segregated in the visual cortex and subsequently integrated into a comprehensive schema within the OFC. By investigating this transition, we aim to uncover the neural architecture that enables the effortless construction of a coherent reality from a limited, ever-changing viewpoint.

## 2. Materials and Methods

### 2.1 Participants

A total of eighteen right-handed students from Peking University (9 females, 9 males; mean age = 20.4 years, range: 18–26) participated in this study. All participants reported no history of psychiatric or neurological disorders. Written informed consent was obtained from all participants prior to the study, and the protocol was approved by the Ethics Committee at the School of Psychological and Cognitive Sciences, Peking University.

### 2.2 Experiment Design and Procedure

#### 2.2.1 Virtual Environment

We developed a three-dimensional virtual environment using Unity (Unity Technologies, San Francisco; version 2020.1.17.f1c1). The environment featured a uniform blue sky, a terrain containing a centrally located sunken arena (240 virtual meters in diameter), and an encircling upward slope that defined the arena boundary. A circular brown platform (4 virtual meters in diameter) was positioned at the center of the arena to mark its midpoint. Within this base layout, two distinct environments—nature and city—were created, each with a unique ground texture and a corresponding set of three landmarks (Fig. 1B). The landmarks occupied three of four predetermined, symmetrically arranged locations outside the arena boundary, with adjacent locations separated by 90° around the arena center. Landmark positions and their spatial relationships remained constant throughout the experiment, forming a static landmark configuration.

**Fig. 1.**
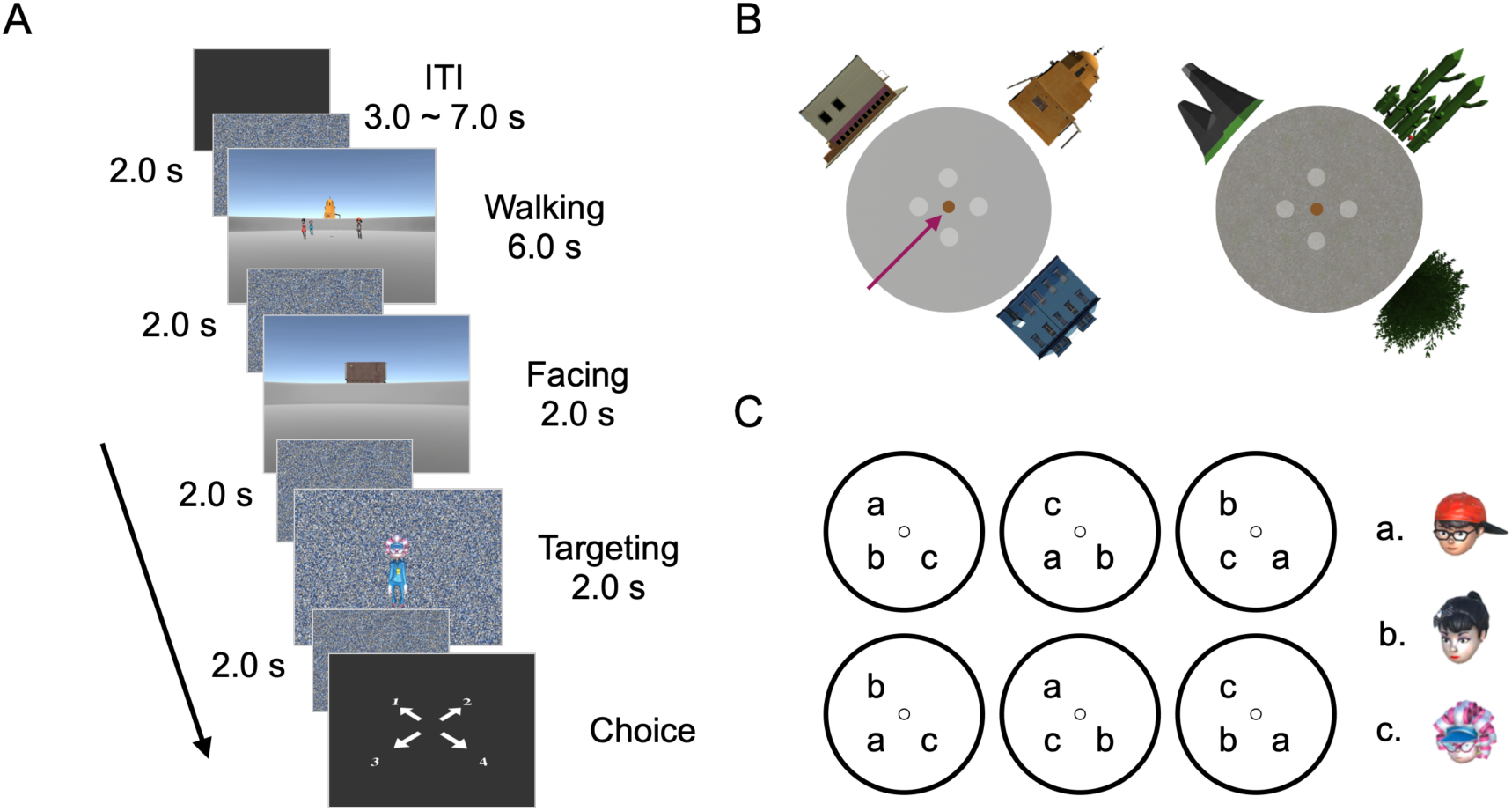
Task design. **A** Automatic space encoding task, example trial. Participants walked toward three human characters and stopped at the center of the arena. They then completed a facing period (orienting toward a character or landmark), followed by a targeting period in which they reported the egocentric direction of a target character or landmark. Task periods were separated by 2.0-s scrambled background screens, and inter-trial intervals ranged from 3.0 to 7.0 s. **B** Two virtual environments. Left: The virtual environment shown in the example trial. The purple arrow indicates the walk-in direction. White dots denote candidate locations for the human characters, and the brown dot indicates the arena center. **C** Human character maps. Of the six maps, two were pseudo-randomly selected for each participant

The task design included two types of spatial elements: landmark sets and human characters. The human characters (Mixamo; https://www.mixamo.com) were pseudo-randomly positioned at three of four vertices surrounding the central platform, with each vertex located 14.15 virtual meters from the center. The spatial arrangement of the human characters defined a “map” (Fig. 1C). The human characters’ positions and their spatial relationships changed across trials, forming a dynamic map configuration.

Based on the two virtual environments and the human character configurations, we designed three behavioral tasks: a voluntary space-encoding (VSE) task, a head-nodding detection (HND) task, and an automatic space-encoding (ASE) task. Participants completed 120 VSE trials (∼60 min) and 160 HND trials (∼45 min) in a behavioral training room on the first day, and performed 144 ASE trials and 16 HND trials, which were pseudo-randomly interleaved (∼80 min in total), during fMRI scanning on the second day. All tasks were conducted from a first-person perspective with a 90° field of view and a 4:3 aspect ratio.

#### 2.2.2 Voluntary Space-Encoding (VSE) task

The task comprised three sessions of increasing difficulty, achieved by progressively increasing the number of additional objects (from one to three; Fig. S1B). The presentation of additional objects was pseudo-randomly balanced. Each trial included three landmarks within the environment, along with additional objects positioned at the predefined locations outside the arena boundary (Fig. S1C). While the positions of the additional objects varied across trials, the landmark positions remained fixed throughout the two-day experiment.

At the beginning of each trial, participants were positioned at the center of the arena in a first-person perspective, allowing them to observe the surrounding environment. During a 7.0 s exploration period, participants could freely adjust their viewing direction using the cursor while viewing three fixed landmarks and one to three additional objects located outside the arena boundary. Following this period, a facing stimulus—either a fixed landmark or an additional object—was presented at the center of the display against the environment background for 2.0 s. Subsequently, a target stimulus (a fixed landmark or an additional object) was presented for 2.0 s on a scrambled background. Participants were instructed to imagine themselves facing the initial stimulus within the environment background and report the egocentric direction of the target stimulus relative to their own position (Fig. S1A). No feedback was provided.

The VSE task comprised three trial types: facing landmark–target landmark (LL), facing landmark–target object (LO), and facing object–target landmark (OL). The proportions of trial types varied across sessions: Session 1 (80% LL, 10% LO, 10% OL), Session 2 (60% LL, 20% LO, 20% OL), and Session 3 (40% LL, 30% LO, 30% OL).

#### 2.2.3 Head-Nodding Detection (HND) task

In the HND task, participants walked from one of four starting locations near the circular boundary toward the human characters and stopped at the central platform. During the walking period, participants viewed three human characters and either a landmark or no landmark positioned behind them. The visual stimuli (i.e., the spatial environment viewed from a first-person perspective) were determined by the combination of the landmark set, human character map, walk-in direction, and the allocentric relationship between the human character map and the landmark set (schema). These factors were pseudo-randomly balanced across the experiment. Of the six human character maps, two were pseudo-randomly assigned to each participant. Participants were not informed about the concepts of the human character map or allocentric schema throughout the task. The walking period lasted 6.0 s, during which each character had a 25% probability of nodding its head at a random time point between the start and end of walking. The probabilities that 0, 1, 2, or 3 characters nodded in a given trial were 50%, 31.875%, 11.25%, and 6.875%, respectively. These proportions were chosen to ensure that each character had an equal probability of nodding across trials.

After the walking period, a photo of one of the characters was presented on a grey background along with a visual instruction prompting participants to indicate whether that character had nodded. Participants responded via key press (Fig. S2A). On 50% of trials, the character shown in the photo had nodded. Feedback was provided after each response, with a green border indicating a correct response and a red border indicating an incorrect response.

#### 2.2.4 Automatic Space-Encoding (ASE) task

Each ASE task began with the same walking period as in the HND task. However, the walking period was followed by the facing period and a targeting period. One human character or one landmark was presented at the center of the display against the environmental background for 2.0 s during the facing period, while an image of a different human character or landmark was presented as the target on a scrambled background for 2.0 s during the targeting period. Each of the three task epochs (walking, facing, and targeting) was followed by a 2.0 s delay, during which a scrambled background was displayed. At the end of each trial, participants reported the egocentric direction of the target relative to their own position using a keyboard response; no feedback was provided (Fig. 1A). After completing 20 practice trials of the ASE task to familiarize themselves with the task sequence, participants performed 144 ASE trials and 16 HND trials during fMRI scanning. Participants were notified that the remuneration depended only on performance in the HND trials, although they were also encouraged to perform the ASE task as best as they could. The experiment comprised 160 trials in total and was divided into four runs. The order of environments was counterbalanced across runs. The two human character maps, four walk-in directions, and four allocentric relationships between human character maps and landmark sets (schema) were pseudo-randomly balanced across the ASE trials.

Participants maintained a correct rate above 80% on HND trials during fMRI scanning, indicating that they attended to head nodding rather than intentionally memorizing the spatial arrangement of the characters (Fig. S2B). Following the scanning session, participants completed a post-scan interview to report the strategies they used to solve the task.

#### 2.2.5 Allocentric Spatial Schema and First-Person Spatial Layouts

The ASE task included two types of spatial structures: allocentric spatial schemas and first-person spatial layouts. Allocentric spatial schemas refer to viewpoint-independent representations of the geometrical relationships between the human character maps and the landmark sets, capturing how the human characters were embedded within the overall environments (Fig. 2B). Across the two environments, four distinct allocentric schemas were defined, each specifying a unique configuration of characters and landmarks. These configurations remained fixed throughout the experiment and were learned implicitly, as participants were never presented with a top-down view.

**Fig. 2.**
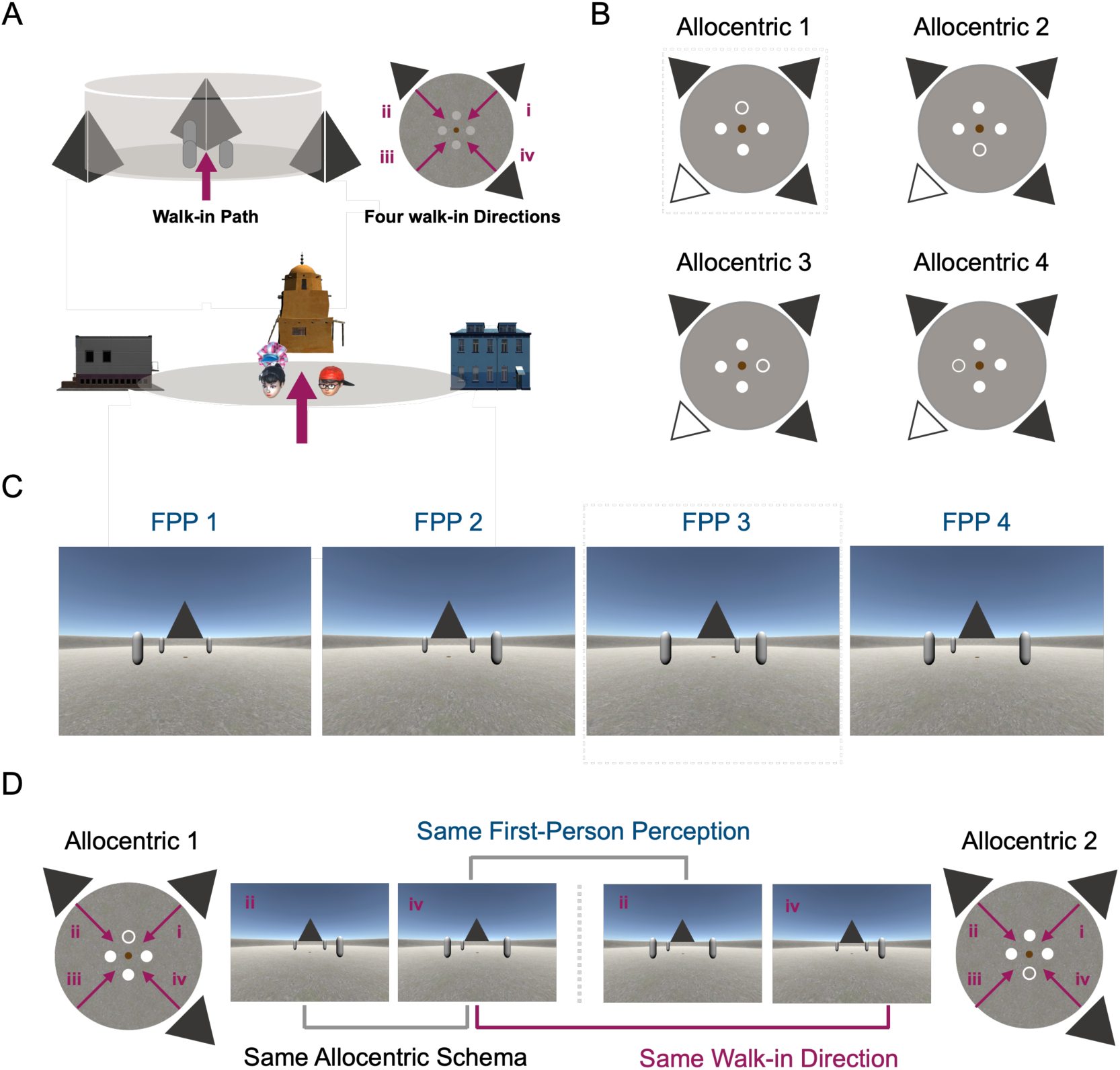
Allocentric spatial schema and first-person spatial layout perception. **A** Schematic illustration of spatial perspectives. Grey cylinders denote human characters, and triangular pyramids denote landmarks. Four walk-in directions are shown in a bird’s-eye view. The spatial layout of human characters relative to the environment can be represented either from a viewpoint-independent allocentric perspective or from an egocentric first-person perspective. Bottom: Spatial layout in the example trial in Fig. 1. **B** Allocentric spatial schemas. Four distinct allocentric schemas depict the spatial relationships among human characters and landmarks. Filled triangles indicate landmark locations, whereas unfilled triangles indicate positions without landmarks. Filled circles denote human character locations, and unfilled circles indicate unoccupied positions. **C** First-person spatial layout perceptions. Four distinct first-person spatial layouts illustrate the egocentric visual configurations perceived during the walking period, which depend jointly on the underlying allocentric schema and the direction from which participants entered the environment. Triangles represent landmarks, and cylinders represent human characters. **D** Relationship between allocentric spatial schemas and first-person spatial layout perceptions. Different first-person spatial layouts can arise from the same allocentric schema, whereas identical first-person spatial layout can emerge from different allocentric schemas

In contrast, first-person spatial layouts describe the egocentric visual organization of characters and landmarks as perceived from the participant’s current viewpoint during navigation. These layouts were jointly determined by the underlying allocentric schema and the direction from which participants entered the environment, yielding four distinct first-person perceptions (FPP; Fig. 2C). Consequently, the same allocentric schema could give rise to different first-person spatial layouts, and identical first-person visual layout could correspond to different allocentric schemas (Fig. 2D; Fig. S4A). Thus, first-person spatial layouts reflect transient, viewpoint-dependent visual experiences, whereas allocentric spatial schemas represent stable, memory-based spatial structures that support interpretation and behavior beyond the immediately visible scene.

### 2.3 Data Acquisition and Analysis

#### 2.3.1 fMRI Data Acquisition

Imaging data were acquired on a 3T Siemens Prisma MRI scanner using a standard 64-channel head coil. Functional images were collected with a gradient-echo echo-planar imaging (EPI) sequence (TR = 2000 ms, TE = 30 ms, flip angle = 90°, 62 slices, voxel size = 2.0 × 2.0 × 2.3 mm³). High-resolution T1-weighted structural images were obtained using a magnetization-prepared rapid acquisition gradient echo (MPRAGE) sequence (TR = 2530 ms, TE = 2.98 ms, flip angle = 7°, 192 slices, voxel size = 0.5 × 0.5 × 1.0 mm³). Head motion was minimized using foam padding placed within the head coil. Visual stimuli were presented via an LCD projector onto a screen positioned above the participant’s head.

#### 2.3.2 fMRI Data Preprocessing

Functional MRI data were preprocessed using FSL FEAT (FMRIB Software Library v6.0)^[24]^. For each session, the first three volumes were discarded to allow for T1 equilibration. The remaining volumes were slice-time corrected, motion-corrected by realignment to the first volume, and temporally high-pass filtered with a cutoff of 100 s. Motion outliers were identified and modeled as nuisance regressors in the first-level analyses. For group-level analyses, functional images from each session were registered to the MNI152 standard space using FSL FLIRT ^[25]^. To preserve fine-grained spatial information for representational similarity analysis, no spatial smoothing was applied.

#### 2.3.3 Representational Similarity Analysis

Task-relevant neural representations were examined using representational similarity analysis (RSA). Neural activity was analyzed separately for the walking (6.0 s), facing (4.0 s), and targeting (4.0 s, including the subsequent delay) periods. For each period, trial-wise multivoxel activity patterns were estimated using a univariate general linear model (GLM), in which BOLD responses for individual trials were modeled with boxcar regressors.

Two classes of nuisance regressors were included in each GLM. The first class was period-specific and modeled the remaining two task periods according to the type of spatial element involved ^[4]^. For example, in the GLM estimating walking-period activity, the facing period was modeled with two nuisance regressors corresponding to the facing element type (static landmark vs. dynamic human character), and the targeting period was modeled with two regressors based on the target element type. This procedure reduced contamination from other task events when estimating trial-specific responses for the period of interest. Accordingly, separate GLMs were constructed for the walking, facing, and targeting periods, each including a distinct set of period-specific nuisance regressors.

The second class of nuisance regressors, shared across all GLMs, included regressors for HND trials, response-related keypress cues, and six head-motion parameters. This modeling procedure yielded trial-wise multivoxel activation patterns for each task period and experimental session in each participant’s native space.

#### 2.3.4 Searchlight-Based RSA

Searchlight-based representational similarity analysis was performed to quantify task-relevant information encoded in local multivoxel activity patterns. For each voxel, a spherical searchlight with a 6 mm radius was defined. Within each sphere, trial-wise multivoxel activation patterns were extracted and organized into a matrix with trials as columns and voxels as rows. Pattern similarity between all pairs of trials was computed using Pearson correlation. Correlation coefficients were assigned to the center voxel of each sphere and transformed using Fisher’s *r*-to-*z* normalization, yielding a trial-by-trial similarity matrix for each task period at each voxel.

To isolate the contribution of task-relevant spatial information while controlling for potential confounds, a second-level GLM was applied to the similarity matrix for each task period. Each categorical regressor encoded whether a trial pair shared the same spatial information (coded as 1) or differed (coded as 0). Separate GLMs were constructed for the walking, facing, and targeting periods, each including regressors corresponding to spatial variables of interest. For the walking period, regressors modeled allocentric spatial schemas, first-person spatial layouts, human character maps, and walk-in directions (with or without a visible landmark). For the facing period, regressors included allocentric spatial schemas, first-person spatial layouts, human character maps, and the category of the facing element (human character vs. landmark). For the targeting period, regressors included allocentric spatial schemas, first-person spatial layouts, human character maps, and the category of the target element (human character vs. landmark).

Parameter estimates for each regressor were assigned to the center voxel of each searchlight, producing whole-brain statistical parametric maps for each spatial variable and task period. These maps were averaged across the four scanning runs for each participant and subsequently normalized to a standard MRI template. Whole-brain statistical analyses were conducted within a gray matter mask ^[26]^. Statistical significance was assessed using permutation-based nonparametric testing implemented in FSL Randomise (5,000 permutations)^[27]^, with a voxel-wise threshold of *p* < 0.001 and a cluster-level threshold of *p* < 0.05.

#### 2.3.5 ROI-Based RSA

To validate the clusters identified in the searchlight analysis—including bilateral superior lateral occipital cortex (LOC), left orbitofrontal cortex (OFC), and left lingual gyrus—we conducted independent region-of-interest (ROI)–based RSA using spherical ROIs. For the LOC, left OFC, and lingual gyrus, a 10-mm radius sphere was used ^[28]^. Because the searchlight-defined cluster in the right lingual gyrus was comparatively smaller, a 6-mm radius sphere was used for this region. For the hippocampus, an anatomical ROI was defined using the Harvard–Oxford subcortical structural atlas. The anterior hippocampus was delineated as voxels anterior to the final slice of the uncus (y = −21)^[29]^ and used as an anatomical mask for small-volume correction. Within this region, a 5-mm radius sphere was centered on the cluster identified by small-volume correction (voxel-level *p* < 0.05, cluster-level *p* < 0.05) to conduct an independent ROI-based RSA.

All ROIs were resampled and aligned to each participant’s native space, and voxels outside the brain were excluded. To control for spatial autocorrelation effects on similarity estimates, a permutation-derived chance level was subtracted from observed similarity values within each ROI. Specifically, trial labels in the correlation matrix were randomly permuted prior to computing pattern similarity for “same” and “different” conditions; this procedure was repeated 5,000 times per ROI to generate a null distribution. Baseline-corrected similarity values were then computed by subtracting the chance level and averaging across the four scanning sessions. Finally, the baseline-corrected similarity values were tested against zero across participants using a two-tailed one-sample *t* test.

#### 2.3.6 Statistical Analysis

For searchlight-based RSA, we used an initial threshold of *p* < 0.001. If no clusters were revealed, a liberal threshold of *p* < 0.01 was used. The reliability of clusters was tested using a nonparametric statistical inference that does not make assumptions about the distribution of the data ^[27]^; the test was conducted with the FSL randomize package (version v6.0.7.12, http://fsl.fmrib.ox.ac.uk/fsl/fslwiki/Randomise) and performed 5000 random sign-flips on whole-brain searchlight beta images; we then reported clusters with the size higher than 95% of the maximal suprathreshold clusters in permutation distribution. For ROI-based analysis, a two-tailed *t*-test was used to examine the RSA. Bonferroni-correction was applied for multiple comparisons. One-way analysis of variance was used to test the behavioral correct rate on each task condition. The statistical significance was determined according to whether the corrected *p* value was smaller than 0.05.

## 3. Results

### 3.1 Behavioral Task and Results

To examine the scene perception including both mobile objects and immobile landmarks, we devised the automatic space-encoding (ASE) task. In the ASE task, each trial began with a walking period, during which participants were instructed to attend to the head nodding of three human characters within one of the two virtual environments (Fig. 1B). Each virtual environment contained three different immobile landmarks, of which only one or none of them was visible during the walking period. Following the walking period, a human character or landmark was presented within the environment, and participants were instructed to imagine themselves facing it. After the facing period, another human character or landmark was presented on a scrambled background as a target stimulus, and participants reported the direction of the target character or landmark relative to their imagined self-position in the environment. The same three human characters and two environments were used throughout the experiment.

To minimize intentional memorization of spatial layouts, we interleaved head-nodding detection (HND), which required the participants to judge the head-nodding of human characters immediately after the walking period (Fig. S2A), and ASE trials pseudo-randomly. Participants were instructed to focus on head-nodding detection during the walking period in all trials. Monetary incentives were tied exclusively to performance on HND trials, and trial type was distinguishable only after the walking period.

On the day before performing the ASE task during scanning, participants were familiarized with the two virtual environments and three human characters through the voluntary space-encoding (VSE) task (Fig. S1A) and the HND task (see Methods).

### 3.2 Basic behavioral results

A total of eighteen participants completed the experiments (40 trials × 4 runs). 90% of trials were ASE trials, and 10% were HND trials. In the HND task, participants exhibited above 80% correct rate for both head-nodding and no-nodding trials (Fig. S2B). In the ASE task, participants exhibited high performance, with a mean correct rate of 89.81% ± 0.79% (mean ± SD, n = 18). Performance did not differ significantly across the two virtual environments (*t* [17] = -1.09, *p* = 0.290, Cohen’s *d* = -0.26), nor across the six human-character maps, as revealed by a one-way ANOVA (*F* [5, 30] = 2.43, *p* = 0.058). In the ASE task, the landmarks occupied three of four predetermined, symmetrically arranged locations outside the arena boundary, leaving one location empty. This arrangement resulted in one walk-in direction in which participants could not see a landmark behind the human characters. We examined whether the landmark visibility influenced participants’ performance and found no significant difference in correct rates between trials with visible versus invisible landmarks (*t* [17] = -0.74, *p* = 0.467, Cohen’s *d* = -0.17; Fig. S4B).

Depending on the spatial elements presented during the facing and targeting period, ASE trials were classified into four types: facing human-targeting human (HH); facing human-targeting landmark (HL); facing landmark-targeting human (LH); facing landmark-targeting landmark (LL). No significant differences in behavioral performance were observed between trials in which the facing and target elements were of the same type (*t* [17] = -1.97, *p* = 0.065, Cohen’s *d* = -0.28), versus different types (*t* [17] = -1.12, *p* = 0.280, Cohen’s *d* = -0.26), indicating that performance was not influenced by specific spatial conditions.

In post-scan interviews, all participants reported that they did not intentionally memorize the spatial arrangement of the human characters or their relationships with landmarks, and none could recall the number of human character maps. Participants consistently reported attending to head-nodding movements during the walking period. Collectively, these behavioral results indicate that the task design effectively enabled the examination of neural representations of spatial elements that were automatically encoded during navigation in scenes containing dynamic human characters and static landmarks.

### 3.3 Lateral Occipital Cortex Encodes Spatial Layout of Human Characters in First-Person Perspective

We first examined brain regions representing spatial layout of human characters in the first-person perspective using searchlight RSA, comparing multivoxel pattern similarity between trial pairs with the same spatial layout and those with different spatial layout. For each participant and run, pattern similarity images were computed in native space, Fisher’s *z*-transformed, projected to MNI space, and averaged across runs. Effects of other task-relevant spatial variables were controlled for in the regression model (see Methods). This analysis revealed significant clusters in the bilateral superior lateral occipital cortex (LOC; voxel-wise *p* < 0.001, cluster-level *p* < 0.05, corrected for multiple comparisons; Fig. 3B). In addition to the LOC, significant clusters were observed in the lingual gyrus (voxel-wise *p* < 0.001, cluster-level *p* < 0.05; Supplementary Table S1).

**Fig. 3.**
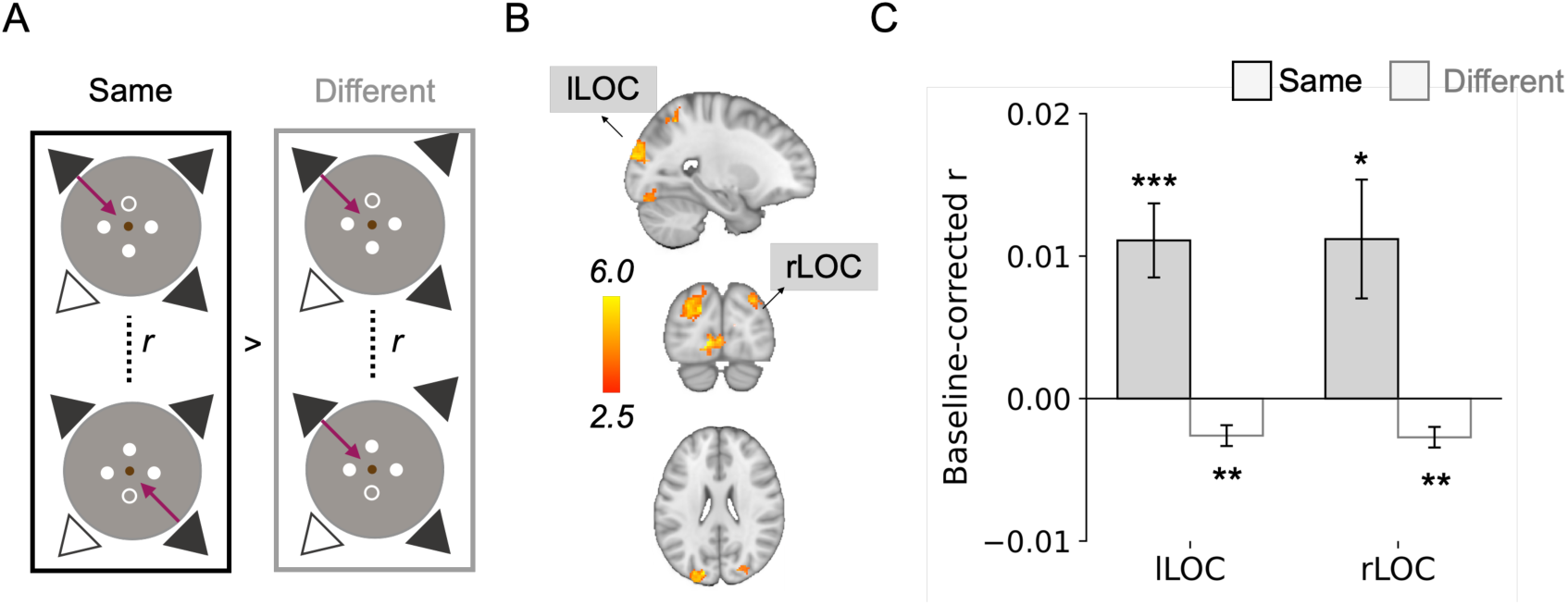
Neural representations of first-person spatial layouts during the walking period. **A** Schematic illustration of decoding first-person spatial layouts using representational similarity analysis (RSA). In the first-person spatial layouts shown on the left and upper right, the lower-left portion was not occupied by a human character, whereas in the lower right allocentric schema, the upper-right portion was not occupied by a human character. **B** Whole-brain searchlight RSA revealed significant clusters in the left and right superior lateral occipital cortex (LOC; voxel-wise *p* < 0.001, cluster-level *p* < 0.05, corrected for multiple comparisons). **C** ROI-based RSA confirmed that multivoxel pattern similarity in bilateral superior LOC was significantly greater than chance for trials sharing the same first-person spatial layout and significantly lower than chance for trials with different layouts (Same: lLOC, *t* [17] = 4.25, *p* = 5.435 x 10^-4^, Cohen’s *d* = 1.00, rLOC, *t* [17] = 2.68, *p* = 0.016, Cohen’s *d* = 0.63; Different: lLOC, *t* [17] = -3.56, *p* = 0.002, Cohen’s *d* = -0.84, rLOC, *t* [17] = -3.74, *p* = 0.002, Cohen’s *d* = -0.88). Significance levels are indicated as *** *p* < 0.001, ** *p* < 0.01, * *p* < 0.05 (Bonferroni-corrected for multiple comparisons; n = 2). Error bars indicate ± SEM

These results were validated using independent ROI-based RSA with spherical ROIs centered on the left and right superior LOC clusters (Fig. 3C). Multivoxel pattern similarity was computed separately for trial pairs with the same versus different spatial layouts and compared against a permutation-derived chance level (5,000 permutations; see Methods). In the left superior LOC, similarity was significantly greater than chance for trials sharing the same spatial layout (*t* [17] = 4.25, *p* = 5.435 x 10^-4^, Cohen’s *d* = 1.00) and significantly lower than chance for trials with different first-person spatial layouts during the walking period (*t* [17] = -3.56, *p* = 0.002, Cohen’s *d* = -0.84). A similar pattern was observed in the right superior LOC, with higher-than-chance similarity for the same-layout condition (*t* [17] = 2.68, *p* = 0.016, Cohen’s *d* = 0.63) and lower-than-chance similarity for the different-layout condition (*t* [17] = -3.74, *p* = 0.002, Cohen’s *d* = -0.88). These findings indicate that the LOC encodes the first-person spatial layouts of the three human characters in the first-person perspective, consistent with previous studies ^[9, 30]^.

### 3.4 Bilateral Lingual Gyrus Encodes the Environment in First-Person Perspective

We next examined brain regions representing the first-person perspective of the environment, as specified by the identity of the currently visible landmark, using searchlight RSA. Trial-wise multivoxel pattern similarity was compared between pairs of trials sharing the same visible landmark identity and pairs with different visible landmarks. This analysis revealed a significant cluster in the bilateral lingual gyrus extending into the occipital fusiform cortex during the walking period (voxel-wise *p* < 0.001, cluster-level *p* < 0.05, corrected for multiple comparisons; Fig. 4B).

**Fig. 4.**
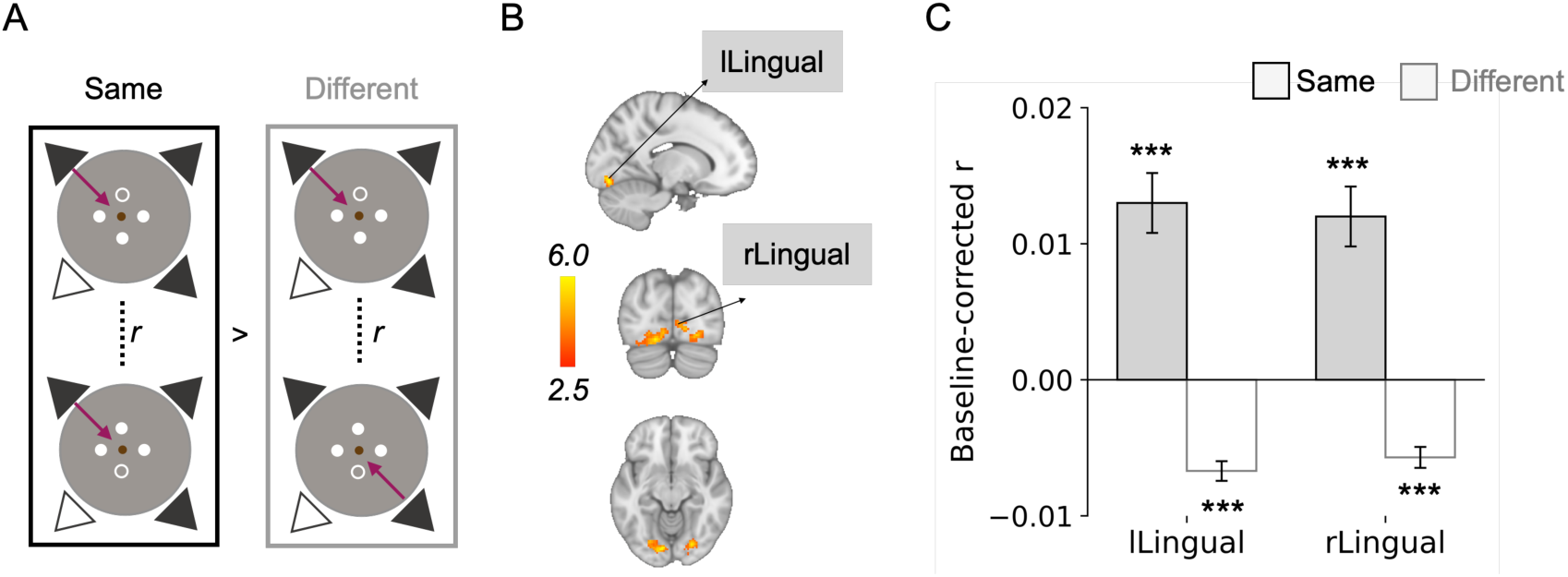
Neural representations of the identity of the viewed landmark during the walking period. **A** Schematic illustration of decoding the identity of the viewed landmark using representational similarity analysis (RSA). In the two trial conditions shown on the left and the upper right, participants viewed the rightmost landmark. In contrast, in the lower right trial condition, participants viewed the leftmost landmark. **B** Whole-brain searchlight RSA identified a significant cluster in the bilateral lingual gyrus and occipital fusiform cortex (voxel-wise *p* < 0.001, cluster-level *p* < 0.05, corrected for multiple comparisons). **C** ROI-based RSA further demonstrated that multivoxel pattern similarity in the left and right lingual gyrus was significantly greater than chance for trials sharing the same task condition (Same: lLingual, *t* [17] = 6.02, *p* = 1.374 x 10^-5^, Cohen’s *d* = 1.42, rLingual, *t* [17] = 5.45, *p* = 4.328 x 10^-5^, Cohen’s *d* = 1.28; Different: lLingual, *t* [17] = -9.27, *p* = 4.666 x 10^-8^, Cohen’s *d* = - 2.18, rLingual, *t* [17] = -7.43, *p* = 9.836 x 10^-7^, Cohen’s *d* = -1.75). *** *p* < 0.001. Error bars indicate ± SEM

The involvement of the bilateral lingual gyrus was further supported by an independent ROI-based RSA (Fig. 4C). In the left lingual gyrus, similarity was significantly greater than chance for trials sharing the same visible landmark (*t* [17] = 6.02, *p* = 1.374 x 10^-5^, Cohen’s *d* = 1.42) and significantly lower than chance for trials with different visible landmarks during the walking period (*t* [17] = -9.27, *p* = 4.666 x 10^-8^, Cohen’s *d* = -2.18). A similar pattern was observed in the right lingual gyrus, with higher-than-chance similarity for the same visible landmark (*t* [17] = 5.45, *p* = 4.328 x 10^-5^, Cohen’s *d* = 1.28) and lower-than-chance similarity for the different visible landmarks (*t* [17] = -7.43, *p* = 9.836 x 10^-7^, Cohen’s *d* = -1.75). These findings indicate that the lingual gyrus and occipitotemporal area encode the first-person perspective of the environment or the identity of the currently visible landmark, consistent with previous studies suggesting that fusiform area activated for identity of attended objects ^[31, 32]^.

### 3.5 Orbitofrontal Cortex Encodes Allocentric Spatial Schemas

Finally, we examined brain regions representing the spatial layout of mobile human characters relative to the environment specified by the immobile landmarks during the walking period. We refer to this space representation as allocentric spatial schema. To identify the brain regions encoding this information, we conducted searchlight RSA comparing multivoxel pattern similarity between trial pairs sharing the same allocentric schema and those with different schemas.

During the walking period, searchlight RSA revealed a significant cluster in the left orbitofrontal cortex (OFC; voxel-wise *p* < 0.001, cluster-level *p* < 0.05, corrected for multiple comparisons; Fig. 5B). This finding was validated by independent ROI-based RSA using spherical ROIs centered on the left OFC (Fig. 5C). In this ROI, similarity was significantly greater than chance for trials sharing the same allocentric spatial schema (*t* [17] = 2.77, *p* = 0.013, Cohen’s *d* = 0.65), whereas similarity for trials with different allocentric schemas did not differ from chance (*t* [17] = -1.55, *p* = 0.139, Cohen’s *d* = -0.37). These results suggest that the OFC encodes allocentric spatial schemas, consistent with previous work implicating OFC in representing abstract, value-based schemas ^[33]^.

**Fig. 5.**
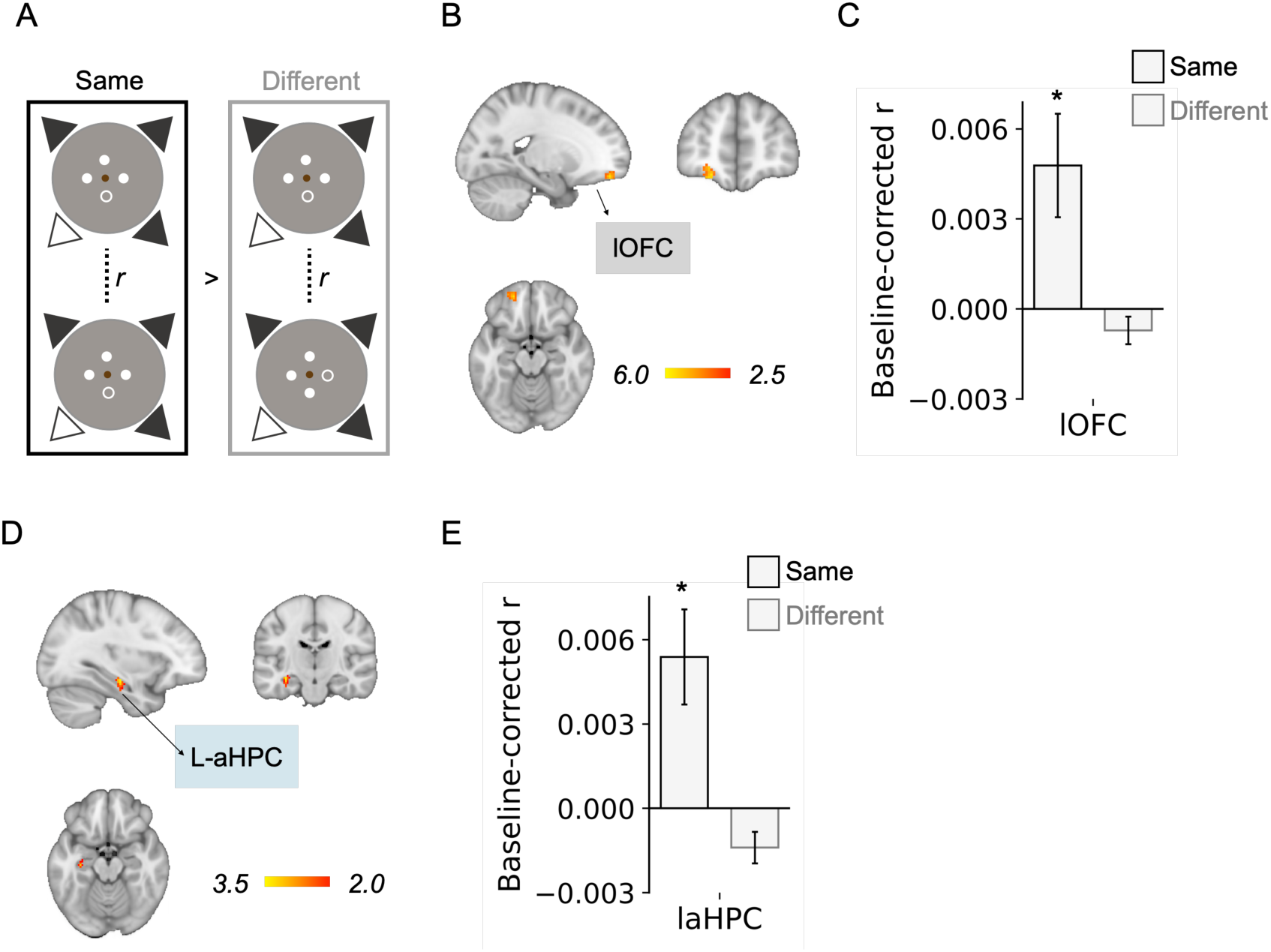
Neural representations of allocentric spatial schemas. **A** Schematic illustration of decoding allocentric spatial schemas using representational similarity analysis (RSA). In the two allocentric schemas shown on the left and the upper right, human character locations were consistently embedded within the landmark configuration, such that the candidate position between the rightmost landmark and the empty location remained unoccupied. In contrast, in the lower right allocentric schema, the unoccupied candidate position was located between the middle and rightmost landmarks. **B** Whole-brain searchlight RSA identified a significant cluster in the left orbitofrontal cortex during the walking period (lOFC; voxel-wise *p* < 0.001, cluster-level *p* < 0.05). **C** ROI-based RSA further demonstrated that multivoxel pattern similarity in the lOFC was significantly greater than chance for trials sharing the same allocentric spatial schema, whereas no significant deviation from chance was observed for trials with different schemas (same: *t* [17] = 2.77, *p* = 0.013, Cohen’s *d* = 0.65; different: *t* [17] = -1.55, *p* = 0.139, Cohen’s *d* = -0.37). **D** Whole-brain searchlight RSA revealed a significant cluster in the left anterior hippocampus (aHPC; voxel-level *p* < 0.05, cluster-level *p* < 0.05 with small volume correction on left anterior hippocampus). **E** ROI-based RSA demonstrated pattern similarity in the left aHPC was significantly higher than chance level for trials sharing the same allocentric spatial schema and not significantly different from chance for trials with different schemas (same condition aHPC, *t* [17] = 2.70, *p* = 0.015, Cohen’s *d* = 0.75; different condition, *t* [17] = 1.76, *p* = 0.097, Cohen’s *d* = -0.59). * *p* < 0.05. Error bars indicate ± SEM

Following the walking period, searchlight RSA showed that allocentric spatial schemas were represented in the left anterior hippocampus (aHPC) during the facing period (voxel-wise *p* < 0.05, small-volume cluster-corrected *p* < 0.05; Fig. 5D), rather than in the OFC (voxel-wise *p* < 0.001, whole-brain cluster-level *p* < 0.05). This effect was confirmed by independent ROI-based RSA: similarity was significantly greater than chance for trial pairs sharing the same allocentric spatial schema (*t* [17] = 2.70, *p* = 0.015, Cohen’s *d* = 0.75), whereas similarity for trials with different allocentric schemas did not differ from chance (*t* [17] = 1.76, *p* = 0.097, Cohen’s *d* = - 0.59; Fig. 5E). These results suggest an involvement of the HPC in self-localization relative to allocentric spatial cognitive map^[4]^.

In addition, we examined whether the ASE task performance was influenced by task conditions related to the RSA analyses. No significant differences in correct rate were induced by the four first-person perspective spatial layouts of human characters, including (*F* [3, 68] = 0.44, *p* = 0.72; Fig. S4C) or excluding trials in which no landmark was visible (*F* [3, 68] = 0.83, *p* = 0.48; Fig. S4D), four allocentric spatial schemas (*F* [3, 68] = 0.35, *p* = 0.79; Fig. S4F), nor four walk-in directions (*F* [3, 68] = 0.42, *p* = 0.74; Fig. S3B).

## 4. Discussion

The present study investigated the neural mechanisms underlying the automatic encoding of complex scenes containing both dynamic human characters and static landmarks. Representational Similarity Analysis (RSA) revealed a clear functional segregation in the representation of distinct scene elements. Specifically, the superior lateral occipital cortex (LOC) encoded the first-person perspective of the spatial layout of mobile human characters, while the lingual gyrus and occipital fusiform cortex represented the environment as specified by the identity of visible landmarks. Crucially, the orbitofrontal cortex (OFC) integrated these components into an allocentric spatial schema, representing the structural relationship between mobile objects and their static environment. Together, these results demonstrate that the brain automatically decomposes perceived scenes into egocentric object layouts and environmental anchors via specialized visual areas before synthesizing them into a high-dimensional allocentric framework within the OFC, even when scene encoding is incidental to the primary task.

We found that the first-person spatial layouts of human characters were represented in the bilateral superior LOC during the walking period. Anatomically, these clusters overlap with the reported coordinates of the occipital place area (OPA), a region known to encode the geometric structure of immediately visible scenes in a viewpoint-dependent manner ^[9, 11, 34]^. While the absence of an independent functional localizer precludes definitive identification of these clusters as the OPA, our findings suggest that this region’s function extends beyond static geometry to include the spatial configuration of mobile objects. This first-person representation in the LOC may facilitate scene understanding by resolving the spatial arrangement of dynamic agents relative to the observer’s viewpoint ^[30]^.

Our RSA results suggest that the lingual gyrus and occipital fusiform cortex encode the first-person perspective of the environment, specified by the identity of the currently visible landmark. While the current RSA procedure cannot definitively distinguish between the identification of a landmark stimulus in isolation and its recognition as a spatial anchor for the egocentric environment, prior evidence supports an integrative role. Ventromedial occipito-temporal regions—including the lingual and fusiform gyri as well as the posterior parahippocampal cortex—have been implicated in landmark-centered representations and recognition of environmentally salient visual cues ^[8, 10, 35]^. Neuropsychological and neuroimaging studies further demonstrate that disruption to these ventral visual regions leads to landmark agnosia and topographical disorientation, characterized by an impaired recognition of orientation-relevant features ^[8, 36]^. Thus, by encoding environmental anchors from a first-person perspective, the lingual gyrus and the occipital fusiform cortex likely enabled the perceptually derived first-person spatial layout to be mapped onto a broader environmental representation extending beyond the current visual field and maintained in memory ^[17]^.

This anchoring process appears to be a prerequisite for the higher-order computations observed in the OFC. While the lingual gyrus and superior LOC decompose the scene into static anchors and mobile object configurations in first-person coordinates, the OFC integrates these distinct streams to construct a comprehensive allocentric spatial schema ^[21]^. This suggests a hierarchical architecture in which the ventral and lateral visual streams provide the necessary spatial "scaffolding"^[37]^ that the OFC then utilizes to maintain a coherent, high-dimensional representation of the scene, even when the encoding is processed incidentally ^[38, 39]^.

The identification of a coherent scene representation in the OFC is consistent with evidence that the orbital prefrontal cortex supports schematic representations that abstract across experiences to facilitate inference ^[20, 40]^. By integrating current perceptual input with memory—rather than relying solely on immediate sensory evidence—these schemas encode invariant relational geometry ^[21, 22]^. Such representations enable the reconstruction of unobserved spatial structures from limited cues ^[41, 42]^, thereby supporting a comprehensive model of the entire scene in allocentric coordinates. The schema identified in the OFC likely differs from the representations reported in the ventromedial prefrontal cortex (vmPFC), which are typically reconstructed from stable and physically observable environmental elements ^[41, 43]^. This distinction suggests that while vmPFC may encode fixed environmental associations, the OFC may be uniquely specialized for synthesizing latent spatial structures when the environment is only partially visible and requires dynamic reconstruction ^[23, 44]^.

Following the initial encoding phase, the left anterior hippocampus (aHPC) exhibited robust allocentric schema representations during the facing period. This finding suggests that once the global scene structure is established, the hippocampus is recruited to localize the self within that abstract spatial framework ^[23]^. This observation aligns with human fMRI evidence showing that the hippocampus anchors the self within a cognitive map ^[4]^ and supports representations that generalize across different environments sharing similar spatial geometry ^[41, 45]^. Converging evidence from macaque electrophysiology further demonstrates the existence of "schema cells" in the anterior hippocampus that maintain consistent firing patterns across distinct environments when neural maps are aligned relative to a hidden goal ^[46]^. Crucially, these hippocampal representations are not driven by immediate gaze direction; instead, they are organized around task-defined states and goal-centered geometry. Together, these findings suggest that the hippocampus integrates current sensory input with memory-based schemas—potentially provided by the OFC ^[23]^—to maintain abstract awareness of one’s position within a structured spatial environment.

Despite these insights, several limitations remain. While our passive paradigm ensured automaticity, future research using active encoding could clarify how top-down attention modulates the OFC-hippocampal circuit. In addition, although RSA identifies key nodes within the scene construction hierarchy, magnetoencephalography or intracranial recordings are needed to resolve the precise directional flow of information between the visual streams, the OFC, and the hippocampus. Nonetheless, this work holds significant implications for our understanding of everyday cognition. It reveals that the brain effortlessly constructs sophisticated mental maps—maintained in allocentric coordinates—by integrating first-person perspectives of both mobile objects (e.g., human characters) and the static environment specified by immobile landmarks without conscious attention. By uncovering the neural scaffolding that bridges immediate perception and abstract memory, we gain a deeper appreciation for how the mind maintains a coherent reality from limited cues.

