## Supplemental File for "The orbitofrontal cortex constructs allocentric schemas by integrating dynamic mobile agents with static environmental anchors"

**Supplementary information**

**Table S1** Brain regions encoding spatial information during the walking period (identified by searchlight-based RSA). For each significant cluster, the peak t value and corresponding MNI coordinates are reported. During the walking period, all listed regions were identified using a voxel-wise threshold of $p < 0.001$ and cluster-level threshold of $p < 0.05$

| First-person spatial layouts (walking period) | | |
|---|---|---|
| t value | MNI coordinate | Anatomy |
| 8.46 | [-10, -86, -4] | Lingual Gyrus |
| 7.44 | [-16, -88, 24] | Lateral Occipital Cortex, Superior division |
| 7.55 | [26, -84, 32] | Lateral Occipital Cortex, Superior division |
| 6.6 | [-22, -64, 56] | Lateral Occipital Cortex, Superior division |

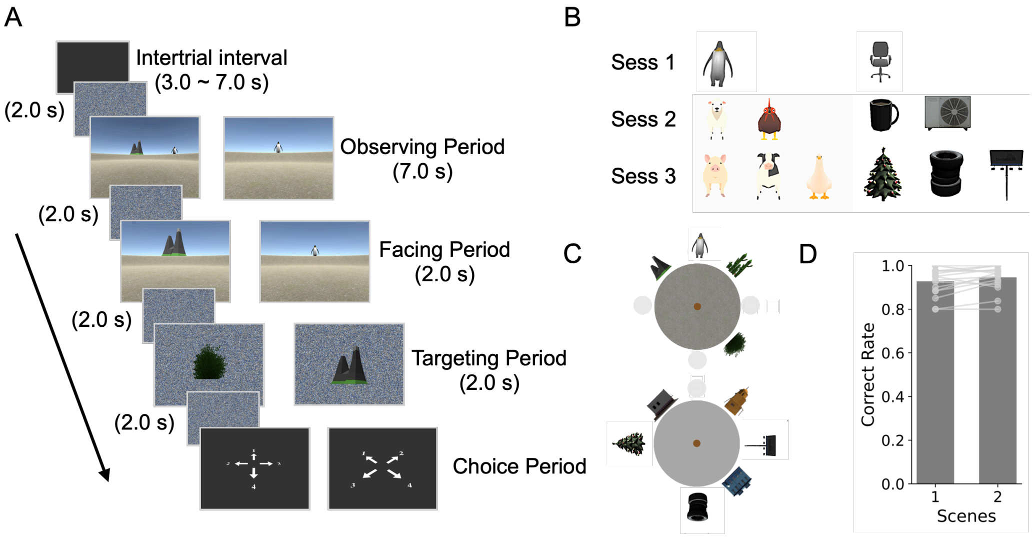


**Fig. S1** Experimental Design of the Voluntary Space Encoding (VSE) task. **A** Schematic of the task structure. Each trial began with a 7.0-s observation period, during which participants were positioned at the center of the virtual arena and allowed to freely scan an environment containing three fixed landmarks and one to three additional dynamic objects. Following the observation period, a "facing element" and a "targeting element" (either a landmark or an additional object) were presented sequentially. Participants were instructed to mentally simulate orienting toward the facing element and to report the egocentric location of the targeting element relative to the imagined heading. **B** Stimulus configuration and progression of task difficulty. To facilitate gradual learning, cognitive load was modulated across training sessions 1-3 by increasing the number of additional objects from one to three. The coordinates of the landmarks remained fixed throughout the experiment, whereas the additional objects were randomly distributed within four designated light-grey circular zones. **C** Depiction of the two virtual environments. These environments used distinct sets of landmarks, identical to those employed in the subsequent automatic space encoding task. **D** Behavioral performance

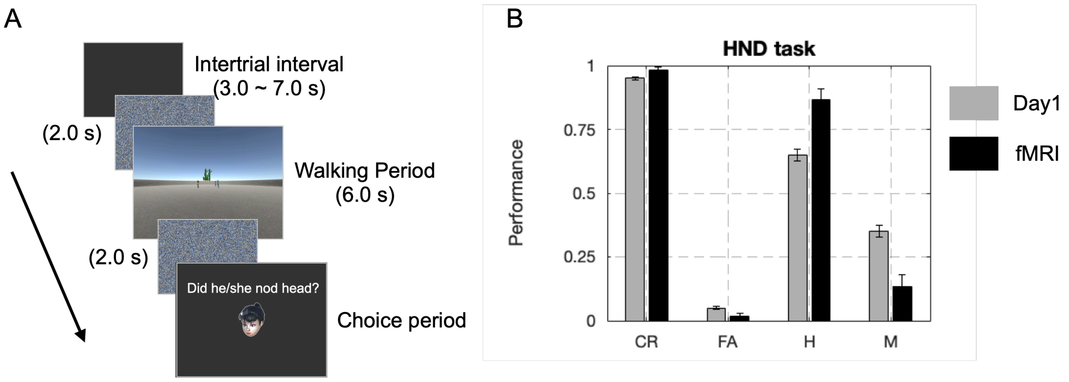


**Fig. S2** Head-nodding detection (HND) task and task performance. **A** Task sequence. During the walking period, participants walked toward three human characters from a first-person perspective and stopped on the central plate. While walking, the three human characters could either nod or remain still at predefined time points. Upon reaching the central plate, participants indicated whether a specific character had nodded by pressing a key. **B** Task performance. In both the practice and fMRI sessions, participants showed a high correct rate in detecting head nodding and high correct rejection rates for trials without head nodding

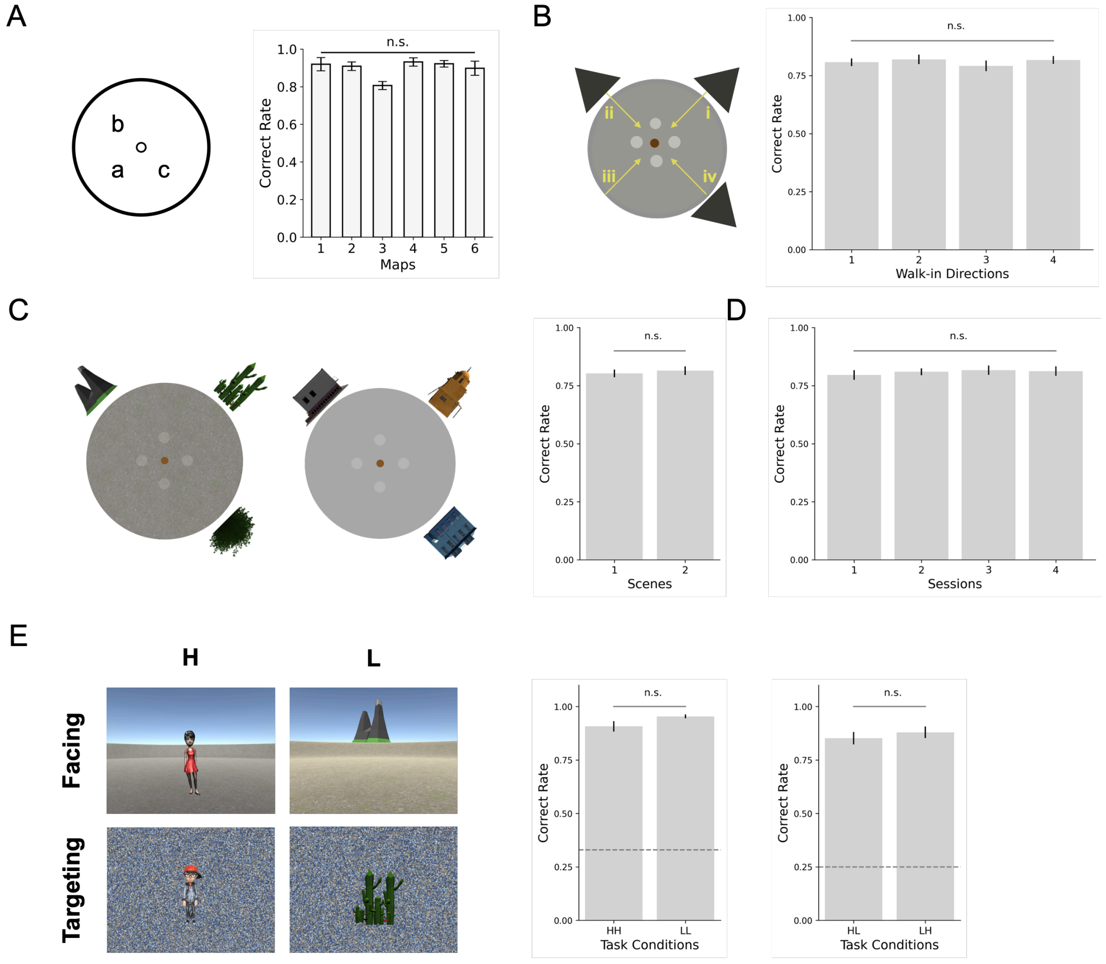


**Fig. S3** ASE task performance across task conditions. **A–D** Control analyses. Retrieval accuracy was not significantly affected by **A** the specific human character map used (*F* [5, 30] = 2.43, *p* = 0.058), **B** the starting direction of the walk (*F* [3, 68] = 0.42, *p* = 0.74), **C** the scene (*t* [17] = -1.09, *p* = 0.290, Cohen's *d* = -0.26), or **D** the experimental session (*F* [3, 68] = 0.22, *p* = 0.88). **E** Retrieval condition. No significant performance differences were observed between trials in which the facing and target elements were of the same type (*t* [17] = -1.97, *p* = 0.065, Cohen's *d* = -0.28) and trials in which they were of different types (*t* [17] = -1.12, *p* = 0.280, Cohen's *d* = -0.26). HH: facing human and targeting human, LL: facing landmark and targeting landmark, HL: facing human and targeting landmark, and LH: facing landmark and targeting human. The dashed line represents chance level

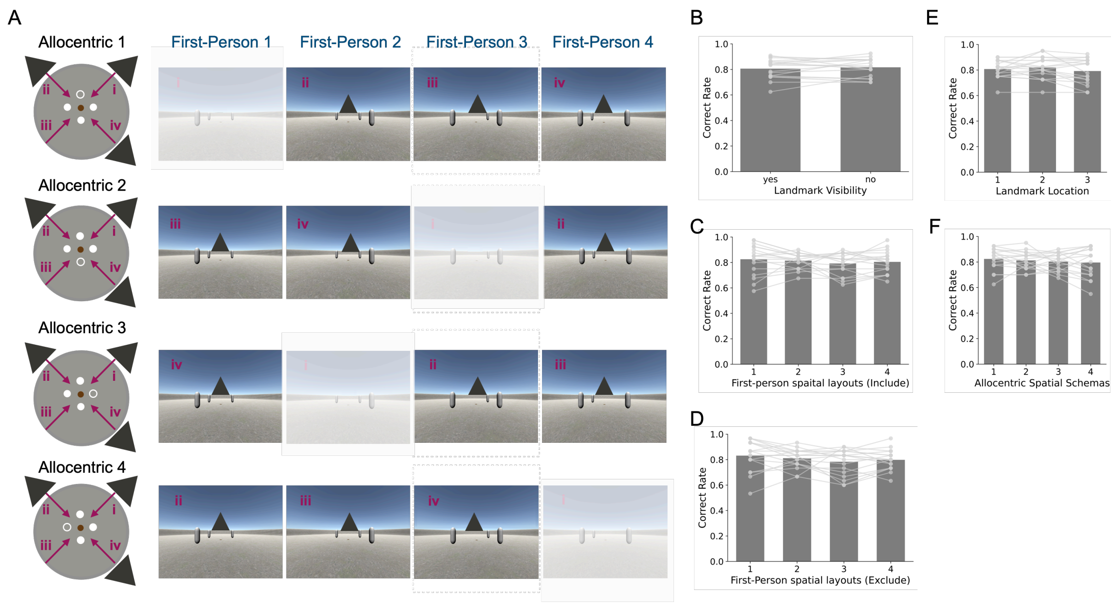


**Fig. S4** ASE task performance across task conditions in relation to RSA. **A** Illustration of how participants' first-person perceptions varied depending on their initial walk-in direction (indicated by purple arrows) within a given allocentric spatial schema. First-person visual perceptions were categorized into four distinct layouts. **B** Participants' performance did not differ significantly based on whether the landmark behind the human characters was visible during the walk ($t$ [17] = -0.74, $p$ = 0.47, Cohen's $d$ = -0.17). **C** Correct rate did not differ significantly among the four first-person spatial layouts when trials without a visible landmark were included ($F$ [3, 68] = 0.44, $p$ = 0.72). **D** Correct rate did not differ significantly among the four first-person spatial layouts when trials without a visible landmark were excluded ($F$ [3, 68] = 0.83, $p$ = 0.48). **E** No significant differences were observed across current viewing landmark locations ($F$ [2, 51] = 0.48, $p$ = 0.62). **F** No significant differences were observed across allocentric spatial schemas ($F$ [3, 68] = 0.35, $p$ = 0.79)

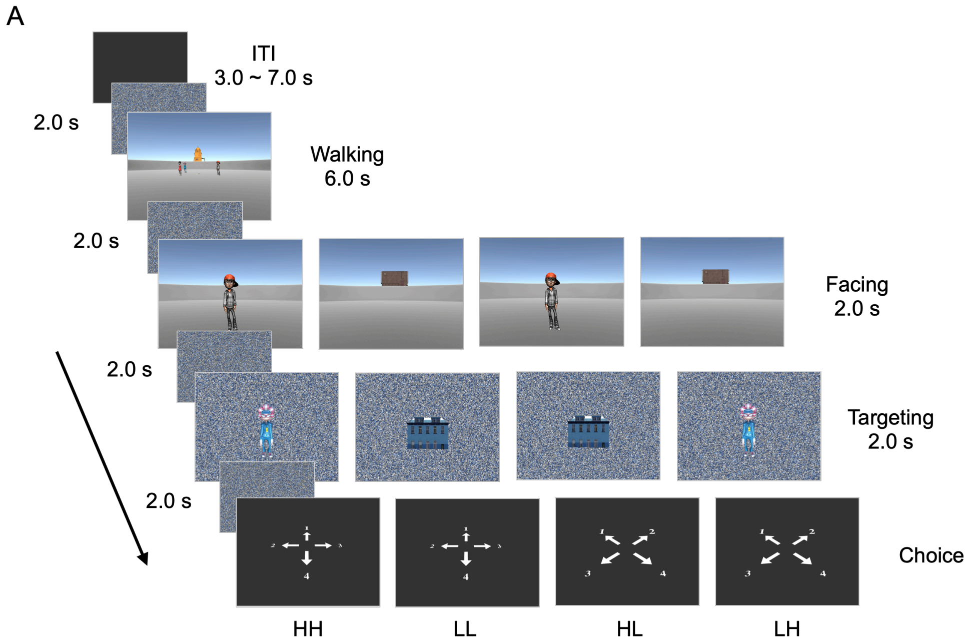


**Fig. S5** Four ASE task conditions. HH: facing human and targeting human; LL: facing landmark and targeting landmark; HL: facing human and targeting landmark; and LH: facing landmark and targeting human
